# Systematics of harvester ants (*Messor*) in Israel based on integrated morphological, genetic, and ecological data

**DOI:** 10.1101/2023.07.16.549226

**Authors:** Maya Saar, Pierre-Andre Eyer, Tali Magory Cohen, Armin Ionescu-Hirsch, Roi Dor, Netta Dorchin

## Abstract

Harvester ants of the genus *Messor* are considered ecosystem engineers, whose distribution is broadly influenced by a variety of environmental factors. Although distinct *Messor* species have been reported to inhabit different habitats, their taxonomy in Israel remains largely ambiguous, hampering the proper ecological characterization of these species. Here, we applied an integrative species delimitation approach combining morphology-based identification, phylogenetic analyses of nuclear and mitochondrial genes, and ecological niche modelling to investigate the phylogenetic relationships among *Messor* species in the small but ecologically diverse region of Israel. Our analyses of mitochondrial genes revealed the presence of at least 13 well-defined lineages, whereas only seven were supported by the analysis of the nuclear genes. However, the concatenated tree that included all the three markers supported 11 lineages. Among two of the lineages-in *M. semirufus* and in a group of ants closest in resemblance to *M. grandinidus-* we identified 3-4 clades that were well established on most trees, inviting further study. In addition, we reveal three undescribed species and raise two subspecies to species rank, highlighting the high diversity of harvester ants in Israel. Ecological niche modelling consistently supported the observed distribution of species, with soil type and average annual temperature being the most influential factors. These results demonstrate that species distribution modelling can serve as a valuable component of integrative species delimitation. We call for future studies to investigate these fascinating lineages of one of the most prominent and ecologically important genera of ants in the Mediterranean Basin.

## INTRODUCTION

Species classification and description represent the first steps to understanding the natural history and ecology of a species (Rojas 1992; Mace 2004; Samper 2004). In addition to these fields of research, species delimitation lies at the nexus of taxonomy and ecology, investigating both micro- and macro-evolution processes to identify the taxonomic boundaries between separately evolving lineages (Bickford et al. 2007; De Queiroz 2007). Ecological niche modelling may supplement systematic studies in accumulating evidence for species delimitation. Different species have been described as different lineages occupying distinct ecological niches (i.e., an adaptive zone minimally different from any other lineages; Van Valen 1976).

Harvester ants are often considered ecosystem engineers, impacting their habitat by preying, distributing, and accumulating plant material, as well as contributing to soil turnover by excavating their underground tunnels (Steinberger et al. 1991, 1992; MacMahon et al. 2000; Jouquet et al. 2006; Azcárate & Peco 2007; Ginzburg et al 2008; Bulot et al. 2016; Mor Mussery & Budovsky 2017; De Almeida et al. 2020a, b; Uhey & Hofstetter 2022). Harvester ants depend to a great extent on physical attributes of their environment, hence their geographic distribution is known to be influenced by a variety of abiotic factors, such as topographic and vegetation gradients, as well as soil type, texture, and water-retention capacity (Johnson 1992; 2000). These abiotic factors primarily influence species distribution ranges in ants by affecting colony foundation and subsequent establishment when colonies are small and fragile (Tschinkel 1992; Kaspari & Vargo 1995; Johnson 1998). Differences in species ranges that reflect variations in abiotic factors have been observed in different harvester ant genera in the Sonoran and Mojave Deserts. For example, *Pogonomyrmex rugosus* Emery 1895 preferentially nests in coarse-textured soils close to mountains, whereas *Veromessor pergandei* Mayr 1886 nests in finer-textured soils away from mountains (Johnson 1992). Differences have also been reported between congeneric species, where *P. rugosus* and *P. barbatus* Smith 1858 differ in their microhabitats, with *P. barbatus* nesting in soils with a higher content of clay and/or higher water-retention (Johnson 2000).

The harvester ant genus *Messor* (Formicidae: Myrmicinae: Pheidolini) comprises approximately 129 species, the majority of which occur in the Mediterranean Basin (Plowes et al. 2012; Bolton 2023). The last global revision of this genus was conducted about 100 years ago (Emery 1922). Since then, only partial revisions, limited to certain geographic regions or specific species groups, have been undertaken (Arnol’di 1977; Barech et al. 2020; Bolton 1982; Collingwood & Agosti 1996; Schlick-Steiner et al. 2006; Steiner et al. 2011; Steiner et al. 2018; Tohmé & Tohmé 1981). In the most recent checklist of the ants of Israel, 24 *Messor* species and subspecies were listed (Vonshak & Ionescu-Hirsch 2009). Systematics of the genus in this area has been based solely on morphological attributes and based on idiosyncratic approaches employed by different taxonomists, resulting in numerous unresolved taxonomic issues. For example, Santschi (1927) split *M. semirufus* into two subspecies: *M. semirufus semirufus* and *M. semirufus grandinidus* with a total of 13 varieties. Eight of these varieties, which occur in Israel, were later raised to species rank by Tohmé & Tohmé (1981), but Kugler (1988) recognized only three of them as valid species (*M. semirufus, M. ebeninus* and *M. dentatus*) and referred to the other five as color varieties without synonymizing them. Overall, the taxonomic status and relationships among Israeli *Messor* species remained largely ambiguous and require further consideration. This situation is further complicated by the fact that the genus is known to include unrecognized cryptic species, as has been reported for other species complexes in Europe (Schlick-Steiner et al. 2006; Steiner et al. 2018). Because *Messor* species are known to establish colonies in different habitats and/or micro-habitats (Baraibar et al. 2011; Díaz 1991; Warburg & Steinberger 1997; Saar et al. 2018), ecological niche modelling may be suitable for differentiating species in this group. This is particularly true in Israel, which constitutes a hotspot for general biodiversity due to the presence of several ecologically diverse regions (Tchernov & Yom-Tov 1988; Medail & Quezel 1999; Myers et al. 2000) and its position at a continental crossroads (Furth 1975; Vonshak & Ionescu-Hirsch 2009). Israel also serves as a good region for a study on the ecological diversity of *Messor* given that it hosts a high proportion of the species found across the Mediterranean Basin.

In the present study, we investigated the phylogeographic relationships among *Messor* species on a small-scaled, but ecologically diverse region. We applied an integrative species delimitation approach combining morphological phylogenetic, and ecological modelling analyses on individual ants collected in 62 localities across Israel. We estimated phylogeographic relationships among recognized and unrecognized *Messor* morpho-species by inferring divergence events from nuclear and mitochondrial sequences and used ecological niche modelling to characterize the niche of genetically inferred species. We highlight the influence of ecological barriers in shaping phylogenetic patterns and discuss how the ecological heterogeneity of this region may act as a catalyst to generate diversity in this ant genus.

## MATERIALS & METHODS

### Collection

Ants were collected from October 2018 through December 2019 at all four seasons, in 62 sites across the various habitats represented in Israel, including desert, sand dunes, Mediterranean woodland, and scrub (Table 1). A maximum of 30 workers were collected from one nest entrance at sites. If a colony had more than one entrance, workers were collected from only one entrance. Workers were placed immediately into 15 ml Falcon tubes filled with 98% ethanol for subsequent morphological and molecular analyses and stored at 4°C in the laboratory. *Aphaenogaster* sp. collected in Tel-Aviv served as an outgroup, given that *Aphaenogaster* is considered a sister genus to *Messor*.

**Table 1:**
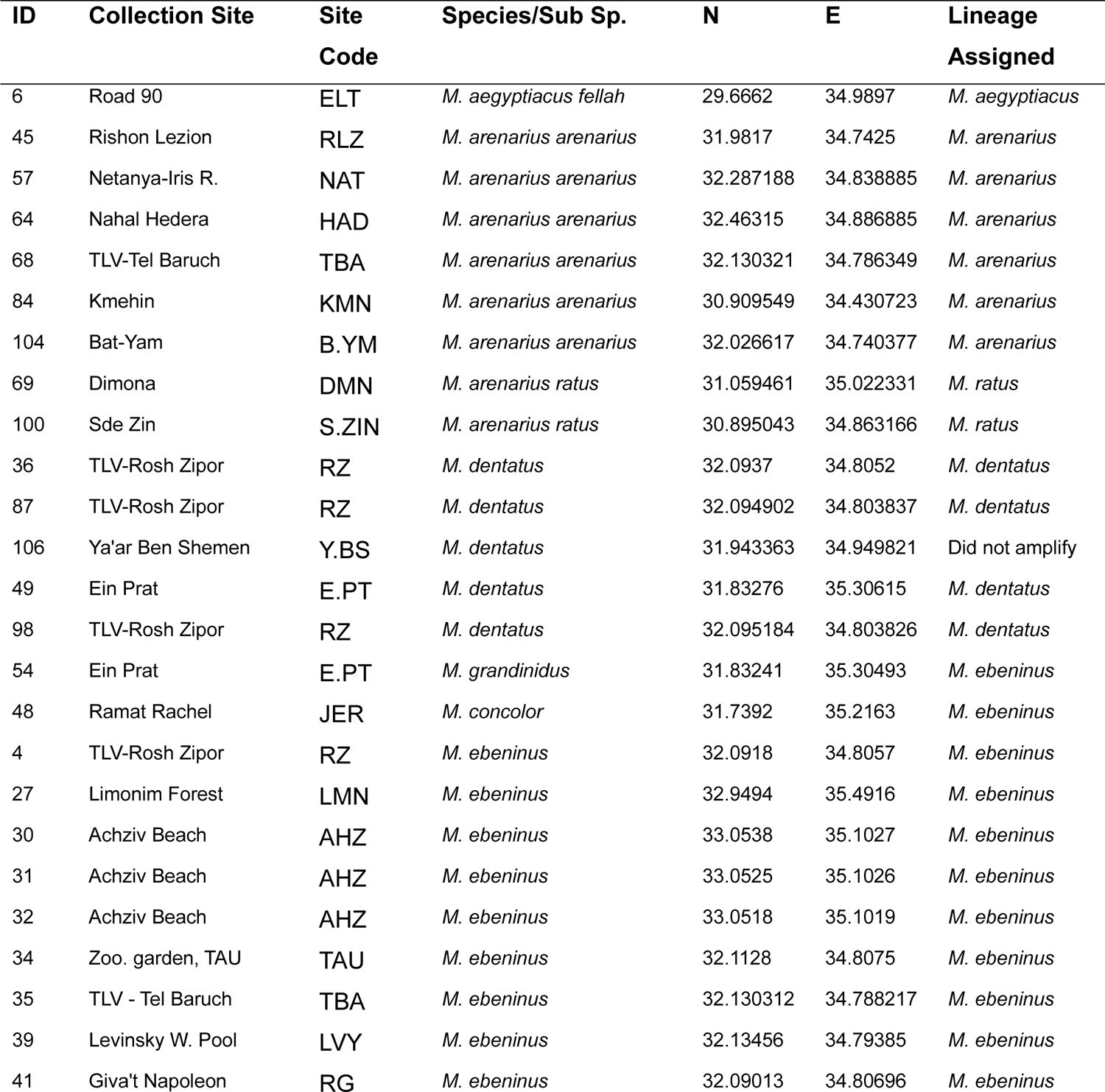
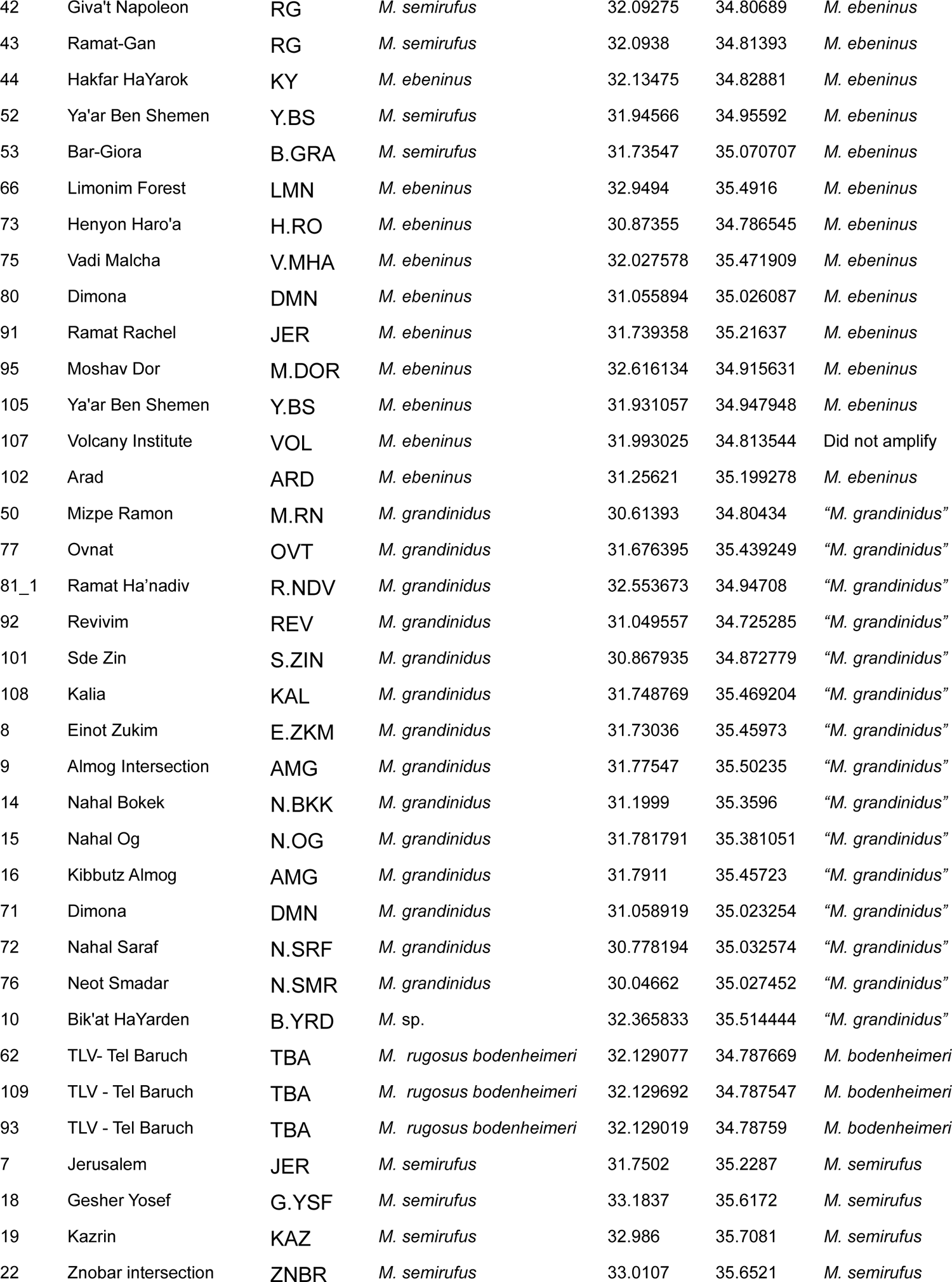
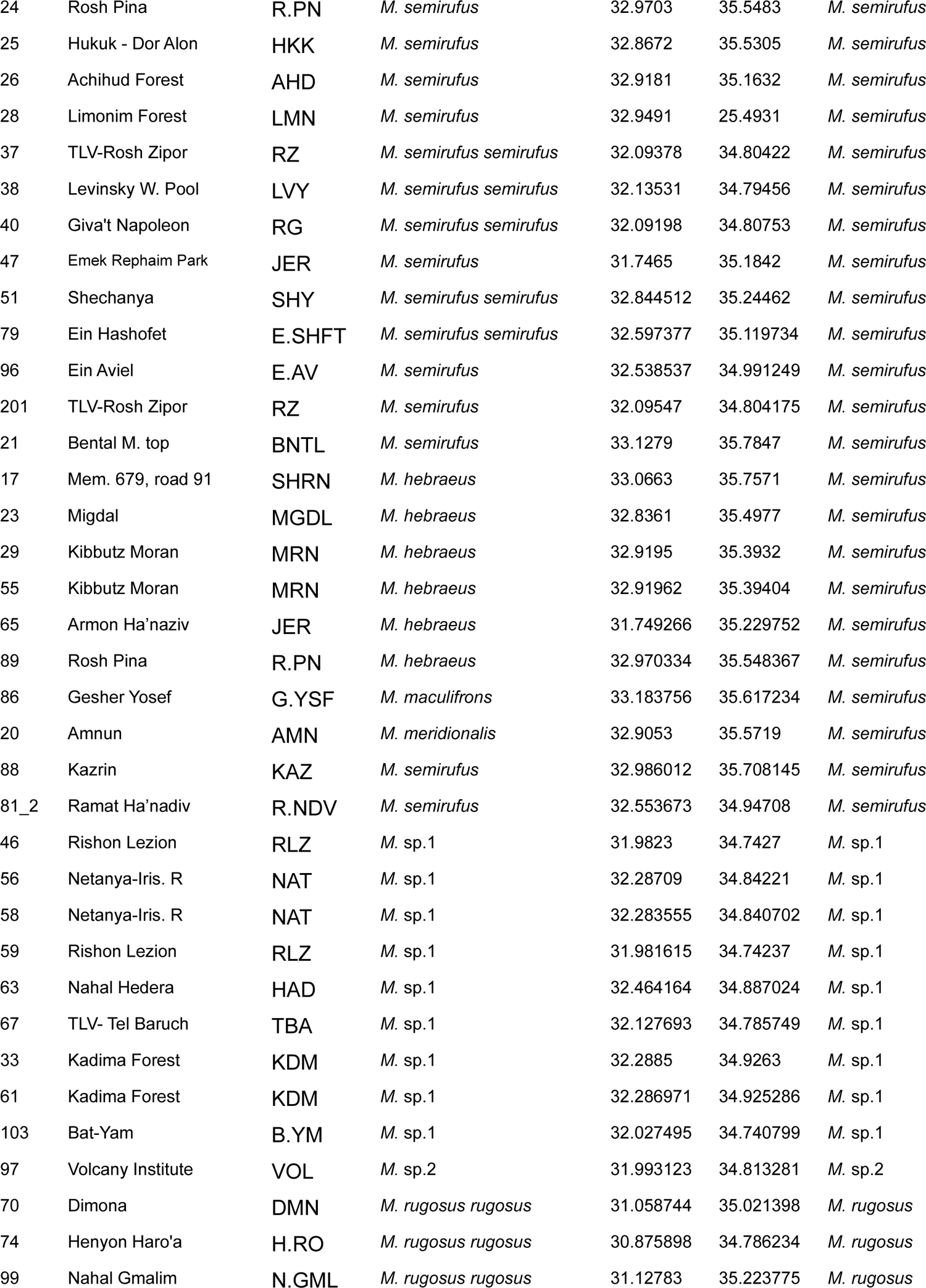
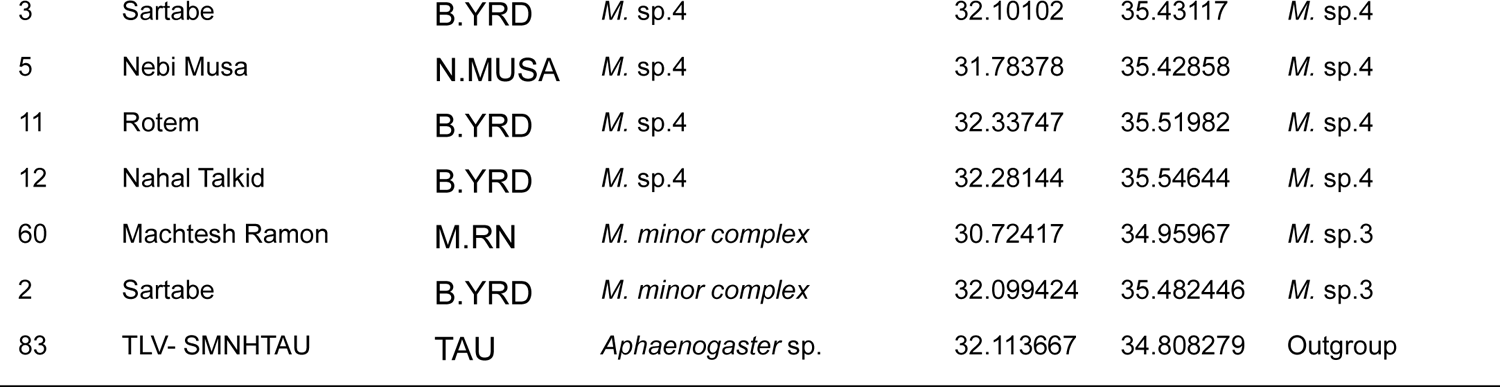
*Messor* samples collected in this study, with information on colony ID, collection site and code, geographic coordinates (N, E), and to which lineage each species/subspecies was assigned to.

### Taxonomy

All individuals were identified to the species level using mainly descriptions by Emery (1908), Santschi (1927), Menozzi (1933), Tohmé (1971), Arnol’di (1977), and Collingwood & Agosti (1996). Individuals that did not key to any known species were labeled *M.* sp., *M.* sp.1, *M.* sp.2, *M.* sp.3 and *M.* sp.4. Also, we named one group of species - “*Messor grandinidus*” because individuals within it were morphologically closest in resemblance to *M. grandinidus* Emery 1912 (Table 1). All specimens collected in this study were deposited in the national collection of insects at the Steinhardt Museum of Natural History, Tel Aviv University, Israel (SMNHTAU).

### Molecular analyses

One individual per colony (overall 102 individuals) was used for DNA extraction. Four legs and the thorax were removed from the ant, comprising 15-25 mg of tissue (depending on ant body size) and were used to extract genomic DNA with the gSYNC^TM^ DNA extraction kit (Geneaid Biotech Ltd.). The remainder of the body of the ant was returned to its specific numbered falcon tube. Nuclear and mitochondrial markers were chosen to infer divergence events across a range of evolutionary timescales. The mitochondrial Cytochrome Oxidase subunit I (COI) was sequenced with the primers *Messorfor2* and *PatMessor* (Steiner et al. 2011), and the mitochondrial Cytochrome B (CytB) was sequenced using the primers *CB1* and *CB2* (Chiotis et al. 2000; Moreau 2008). The nuclear gene Long-wavelength Rhodopsin (LwRh) was sequenced using the primers *LR143F* and *LR639R* (Ward & Downie 2005; Brady et al. 2006). PCR amplification conditions for each marker are detailed in the supplementary material (Tables S1 and S2). Sequencing was performed on an ABI 3730 Genetic Analyzer (Applied Biosystems). Base calling and sequence editing were performed using CodonCode Aligner (CodonCode Corporation, Dedham, MA, USA). For the nuclear marker, reconstruction of parental haplotypes in diploid heterozygous individuals was done using PHASE (Stephens et al. 2001); the PHASE input files were built using SeqPHASE (Flot 2010). Sequences were aligned using the MUSCLE algorithm (Edgar 2004) implemented in CodonCode Aligner.

### Phylogenetic analyses

Bayesian Inference (BI) was used to reconstruct two main phylogenetic trees: One based on the COI mitochondrial marker, and a second tree based on a concatenated dataset combining three markers-two mitochondrial and one nuclear (i.e., COI, CytB, and LwRh). We also constructed three more phylogenetic trees; One tree based on the CytB mitochondrial marker, which we expected to be very similar to the COI tree. A second tree based on the LwRh nuclear marker, and a third tree based on the COI marker which included all samples from the present study as well as related global *Messor* species available from GenBank. Best partitioning schemes and substitution models were calculated using PartitionFinder v.1.0.1 (Lanfear et al. 2012), allocating each codon position of each gene its own substitution model. The best-fitting nucleotide model was selected for each gene using the minimal value of the Bayesian information criterion (BIC). BI was performed with the program MrBayes v.3.2 (Ronquist et al. 2012) under the substitution models estimated with PartitionFinder. Bayesian posterior probabilities were calculated using a Metropolis-coupled Markov chain Monte Carlo approach, with two simultaneous runs starting with random trees. BI was run for 1 × 10^6^ generations, saving one tree every 100 generations to produce 10,000 trees. Convergence and appropriate sampling were ensured by examining the standard deviation of the split frequencies between the two runs (< 0.01) and the potential scale reduction factor diagnostic (PSRF). The first 2,500 trees of each run were discarded as burn-in, and a majority-rule consensus tree was generated from the remaining trees. Trees were constructed in FigTree v1.3.1 (available at http://tree.bio.ed.ac.uk/software/figtree).

For both mitochondrial and nuclear markers, phylogeographic relationships between haplotypes were also represented as networks produced by the median-joining method implemented in NETWORK v.10.1 (Bandelt et al. 1999). For the nuclear LwRh marker, the Haploweb approach was used to define species on the haplotype network. This method identifies independently evolving lineages among samples using diploid phased nuclear markers. The Haploweb approach links haplotypes (*i.e.,* alleles) that co-occur within heterozygous individuals (Flot et al. 2010), and therefore identifies pools of alleles connected through individuals sharing common alleles in their heterozygous state. These pools of alleles denote fields for recombination thus likely represent distinct putative species (Doyle 1995).

### Species Distribution Models (SDM)

To describe the distribution of different lineages identified by the genetic analyses, a set of 33 environmental variables of potential biological importance were selected (Table S3; Steiner et al. 2008). Variables included 19 temperature and precipitation co-factors (CHELSA Climate v.1.1, Karger et al. 2017), nine soil variables (https://soilgrids.org, Hengl et al. 2014, 2017; Itkin et al. 2018), and five remote-sensing-derived vegetation land cover products (Tuanmu & Jetz 2014). All raster data were resampled according to the highest spatial resolution (cell size 0.008) to standardize the model sampling and the individual values of every predictor variable were extracted for each colony and background point. The extent of the boundaries of the state of Israel (34 – 36°N and 29.5 – 33.5°E) was used. The final predictor set was selected using Pearson’s correlation test (Pearson’s |r| <0.8; Table S4), which included 13 variables: soil organic carbon content (stock) [‰ (g kg−1)], coarse fragments [volumetric %], sand [weight %], deciduous broadleaf trees [unit], evergreen/deciduous conifer trees [unit], herbaceous vegetation [unit], mixed/other trees [unit], shrubs [unit], soils category, annual mean temperature [°C], annual temperature range [°C], annual precipitation [mm], and precipitation seasonality [coefficient of variation * 100].

Each lineage identified by the genetic analyses was modeled separately. To minimize sampling bias caused by spatial auto-correlation, a filter to the occurrence records was applied (Boria et al. 2014) with the minimal distance between colonies being at least five km (Steiner et al. 2008). We could not apply the same filtering to the pseudo-absence (background) points because this resulted in very low numbers (∼240 data points). Therefore, background points were generated randomly within the model extent (n = 4001). MaxEnt software package v.3.3.3 (Phillips et al. 2006) was used, which performs well with small sample sizes (Guisan et al. 2007; Pearson et al. 2007), with a regularization multiplier of one and all feature classes to construct species distribution models with clamping. Bootstrapping was applied as the estimation method of error rate via 100 replicates per model. The final datasets included 14 sampling points for “*M. grandinidus”*, 21 for the *M. semirufus*, 25 for the *M. ebeninus* sensu Tohmé, 8 for the *M. arenarius*, and 7 for *M.* sp.1. All other lineages were excluded from the species distribution analyses due to small numbers of sampling points. We estimated the model’s performance of test data by utilizing the threshold-independent value of the ‘Area Under the Curve’ (hereafter AUC) of the Receiver Operating Characteristic. In addition, we generated prediction maps using the model average raw output to identify areas with suitable habitats for each lineage.

## RESULTS

### Species delimitation

A 677 bp fragment of the COI gene was sequenced for 102 individuals, resulting in 70 unique haplotypes. A 403 bp fragment of the mitochondrial CytB was sequenced for 93 of those individuals, resulting in 59 unique haplotypes. The phylogenies generated from the two mitochondrial markers were generally congruent (Figures 1 & S1). After splitting the heterozygous alleles for LwRh, the nuclear dataset contained 96 sequences and 28 unique haplotypes (Figure S2). Both BI and Haploweb analyses revealed the occurrence of different lineages among *Messor* ants collected in Israel.

**Figure 1:**
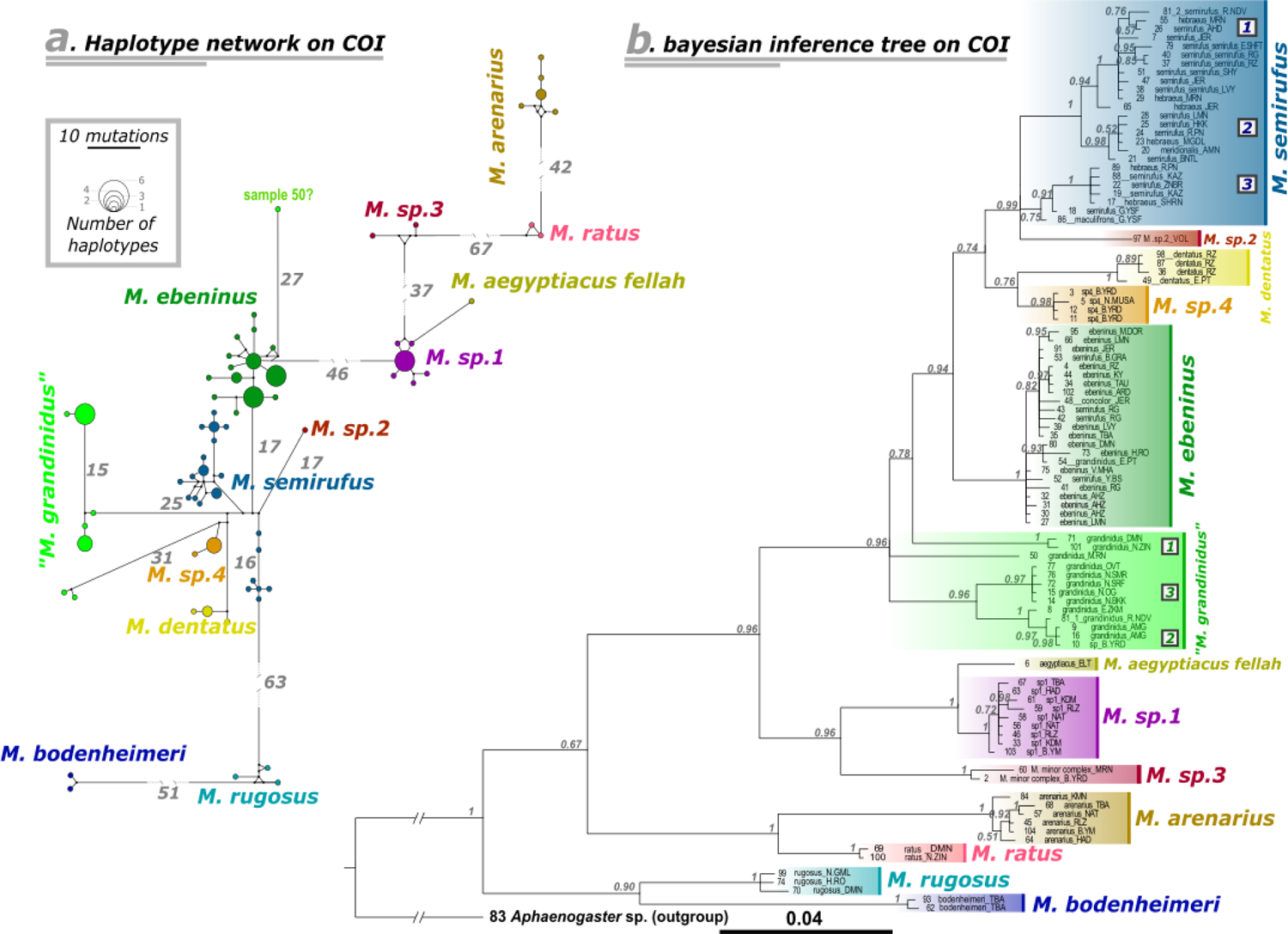
(a) Network of mitochondrial COI haplotypes of *Messor* harvester ants in Israel. Colors correspond to lineages. Branch lengths indicate the number of mutations between haplotypes; circle sizes are proportional to the number of haplotypes observed in the dataset; (b) Bayesian tree of *Messor* in Israel based on the mitochondrial COI marker. Colors correspond to lineages, and branch support values are Bayesian posterior probabilities. Numbers 1-3 in squares represent postulated clades within the *M. semirufus* and “*M. grandinidus*” lineages.

The mitochondrial data suggest that the collected *Messor* in Israel belong to at least 13 distinct species (Figure 1 & S1), while in two of them we identified 3-4 clades within. The concatenated tree supported 11 distinct species out of the 13 (*M. rugosus bodenheimeri* and *M. aegyptiacus fellah* did not amplify on all three markers, Figure 2). Seven of the 13 species recovered were also supported in the analysis of the nuclear marker (Figure S2). With some exceptions detailed below, most of the species supported by the nuclear marker were congruent with those supported by the mitochondrial markers. These results were also consistent when additional sequences from GenBank were integrated in the COI mitochondrial analysis (Figure S3). The following synopsis includes the 13 species of *Messor* found in Israel that were retrieved as distinct genetic lineages in our analyses, with data on their distribution in the country (Figure 3h). The species names used here are derived from the most common morphospecies within each lineage based on the current taxonomic identification of *Messor*.

**Figure 2:**
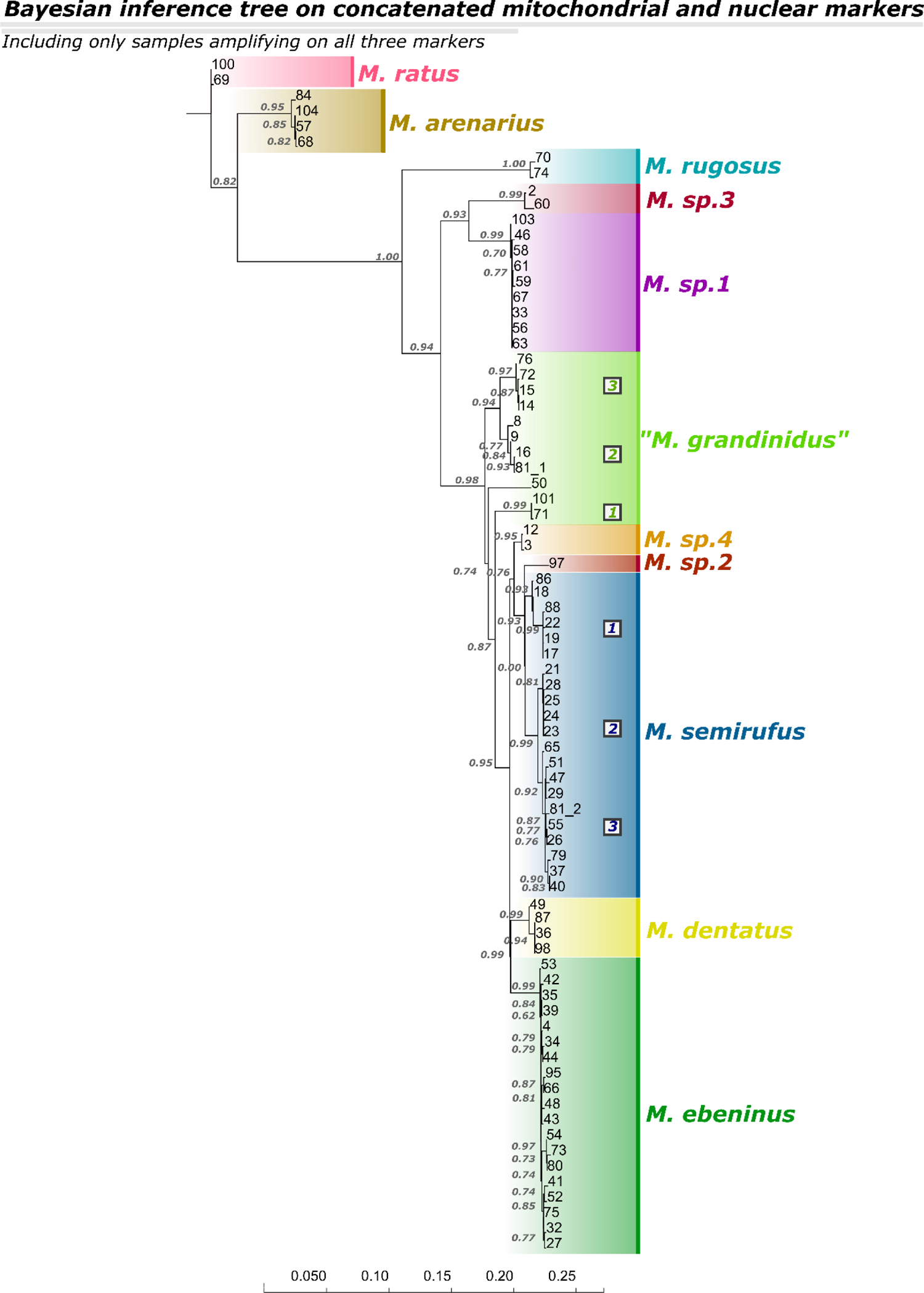
Bayesian tree of *Messor* harvester ants in Israel based on a concatenated genetic dataset. Branch support values are Bayesian probabilities. Numbers 1-3 in squares represent postulated clades within the *M. semirufus* and “*M. grandinidus*” lineages.

**Figure 3:**
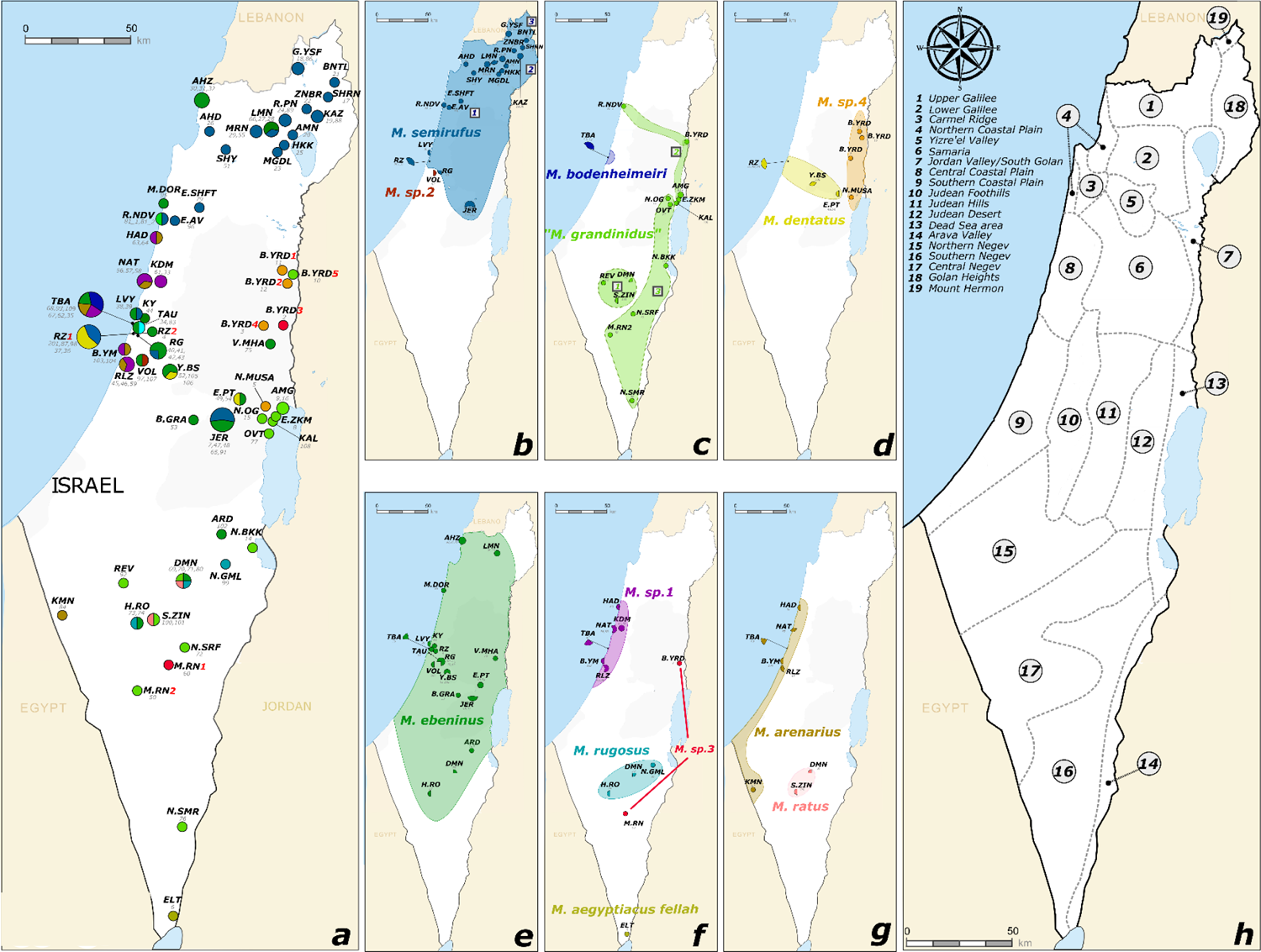
(a) Geographic distribution map of *Messor* harvester ants that were collected in Israel during this study, according to the mitochondrial information. The circle in the lightest blue (TAU - sample 83) represents the outgroup; (b-g) Geographic distribution maps per lineage of *Messor* (Figure 3a un-assembled). In figures (a-g), circle colors correspond to mitochondrial lineages, circle size represents the number of individuals sampled at each locality, the black letters in bold indicate the locality code name, and the gray numbers in Italic indicate the ID of colonies (see Table 1). In (a), the red numbers represent a few localities with the same code name, but different individuals were collected (B.YRD1, B.YRD2 etc.). In (b) and (c), the numbers 1-3 in squares represent different postulated clades within lineages; (h) Illustrated 19 geographic regions within Israel (by Fishelson 1985) that we refer to throughout the manuscript.

### Synopsis of the Messor species in Israel

*Messor arenarius* Fabricius 1787, *Messor ratus* Menozzi 1933 stat. n. Our genetic analyses revealed the presence of two genetic lineages among the samples morphologically assigned to *M. arenarius*, corresponding to the currently recognized subspecies *M. arenarius arenarius* and *M. arenarius ratus*. These two taxa differed substantially from all other lineages for all analyzed markers and consistently formed a monophyletic group (Figures 1, 2, S1, S2, S3). These subspecies are clearly distinguished by genetic data, whereby *M. arenarius arenarius* is distributed in most of central and southern Israel (see a photo of ants; Figure 4c), whereas the Israeli endemic *M. arenarius ratus* is found in the Judean Foothills, Judean Desert and in the Northern and Central Negev (Vonshak & Ionescu-Hirsch 2009, and for our sampling see Figure 3g). Based on these results, we hereby raise *Messor arenarius ratus* to species rank - namely *Messor ratus* Menozzi 1933 stat. n.

**Figure 4:**
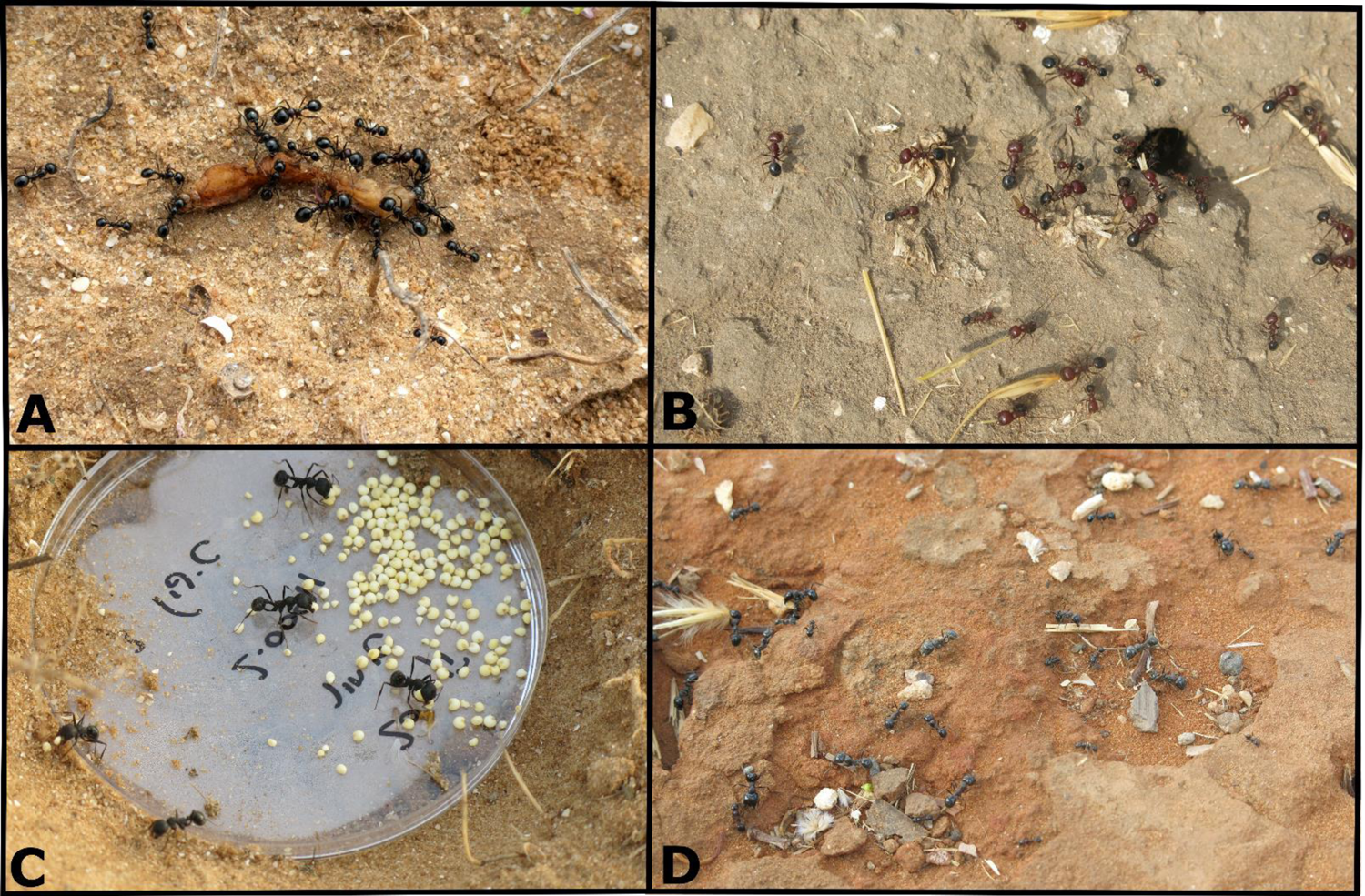
High resolution photos of four lineages of *Messor* harvester ants in Israel: (a) *M.* sp.1 in Tel Baruch; (b) *M. semirufus* in Ramat-Gan; (c) *M. arenarius* in Tel Baruch; and (d) *M. ebeninus* in Ramat-Gan.

*Messor rugosus* André 1881, *Messor bodenheimeri* Menozzi 1933 stat. n. Our analyses revealed the occurrence of two distinct lineages within *M. rugosus,* corresponding to the subspecies *M. rugosus rugosus* and *M. rugosus bodenheimeri,* (Figures 1, S1, S2, S3). This segregation was clearly supported by mitochondrial and nuclear markers as well as by the geographic distribution of these two entities on sandy habitats in Israel, with *M. rugosus rugosus* in the Southern Coastal Plain, Northern and Central Negev, and *M. rugosus bodenheimeri* along the Northern and Central Coastal Plain, and the Carmel Ridge (Vonshak & Ionescu-Hirsch 2009 and for our sampling see Figure 3c, f). Hence, we hereby elevate *M. rugosus bodenheimeri* to species level, namely *M. bodenheimeri* Menozzi 1933 stat. n.

### *Messor ebeninus* sensu Tohmé 1971

*M. ebeninus* formed a distinct monophyletic lineage based on both mitochondrial markers and our concatenated analysis (Figures 1, 2, S1), which, despite its wide geographic distribution (Figure 3e), exhibited low mitochondrial diversity. These results strongly support its status as a distinct species, despite the fact that the nuclear marker did not distinguish between it and *M. semirufus, “M. grandinidus”, M. dentatus*, and *M.* sp.2 (Figure S2). The distribution of *M. ebeninus* includes the entire central region of Israel, spanning from the Upper Galilee in the north and extending southwards to the Central Negev and Arava Valley, making it the most widespread *Messor* species in Israel (Figure 3e, and see a photo of ants; Figure 4d).

### “Messor grandinidus”

We stress that to our knowledge, individuals keyed morphologically to “*M. grandinidus”* in this study are not *M. grandinidus* Emery 1912, but likely related species. We named them after the closest morphological entity. These Individuals formed a paraphyletic group comprising 3-4 clades in all trees (Figures 1, 2, S1, S2, S3). Interestingly, three clades were clearly geographically separated from each other, supporting their status as distinct species. “*M.* grandinidus” occurs along the Jordan Valley through the Dead-Sea area, to the Central and Southern Negev, and the Arava valley (Figure 3c).

### Messor semirufus André 1883

Based on both the mitochondrial markers and our concatenated tree, all the samples morphologically identified as belonging to the *Messor semirufus* complex formed a paraphyletic lineage comprising 3-4 clades (Figures 1, 2, S1, S3, and see a photo of ants; Figure 4b). It seems that the clades in the *M. semirufus* complex are more tied and interconnected, which implies higher gene flow, in comparison with the 3-4 clades within “*M. grandinidus”*. We recommend further extensive sampling to deepen our understanding of the *M. semirufu*s complex and resolve the number of entities that it may possess. In contrast to *M. ebeninus*, *M. semirufus* showed a substantial level of mitochondrial diversity among geographically distant individuals, although the geographic distances among samples was similar between the two species (Figures 3b & e). The mitochondrial genetic diversity found in *M. semirufus* seems to correlate with geographic variation, with samples from northern Israel clustering separately from individuals from other localities. Similar to *M. ebeninus*, *M. semirufus* was also not fully differentiated in the analysis of the nuclear marker. Nevertheless, the clear segregation between *M. semirufus* and other *Messor* species based on both mitochondrial markers, and with comparison to other global *Messor* species-suggests that this lineage is distinct. The distribution of *M. semirufus* in Israel spans from the Golan Heights in northern Israel to Jerusalem (on the Judean Hills), and although it is not clear-cut, mostly-the clades within it correspond to different geographical areas (Figure 3b). *M. semirufus* is probably the second most common complex of species in Israel, following *M. ebeninus*-in terms of distribution.

### Messor dentatus Santschi 1927

Individuals identified as *M. dentatus* formed one distinct lineage in our analyses, that seems distantly related to *M.* sp.4 on both the mitochondrial trees (Figures 1, S1). *M. dentatus* also shared haplotypes with both *M. semirufus* and *M.* sp.4 on the nuclear tree (Figure S2). However, we deduce that *M. dentatus* remains largely distinct as can be seen in all trees and especially in our concatenated tree and when samples from GenBank were integrated in the COI analysis (Figures 2, S3). There seems to be a continuous geographic distribution of entities identified as *M. dentatus*; from the Central Coastal Plain to the Judean Hills. Sample 106 did not amplify on any marker, but it is seen in the geographic distribution, contributing to this continuity (Figure 3d).

### Messor aegyptiacus fellah Santschi 1923

On both the mitochondrial markers, *M. aegyptiacus fellah* formed a distinct lineage (Figures 1, S1, S3). But on the nuclear marker it shared haplotypes with *M.* sp.1, and although morphologically it is clear that these are two distinct species, the nuclear marker analysis supports their close genetic relatedness (Figure S2). We obtained only one specimen of *M. aegyptiacus fellah* from an area close to Eilat (in Arava Valley, see Figure 3f). This subspecies that is endemic to Egypt and Israel was collected in Israel in the past in the Southern Negev and the Arava Valley (Vonshak & Ionescu-Hirsch 2009). The distribution of its higher rank-*M. aegyptiacus-* is known from Israel, Egypt, and North Africa in desert regions (El Bokl et al. 2015; Kugler 1989).

### Additional undescribed species

Our analyses revealed that *Messor* in Israel includes additional, genetically distinct species that are yet to be described. One of these, *M.* sp.1 (see a photo of ants; Figure 4a), found along Israel’s Central Coastal Plain (Figure 3f), was consistently supported in both the mitochondrial, nuclear, and concatenated analyses (Figures 1, 2, S1, S2, S3). Despite its genetic relatedness to *M. aegyptiacus fellah, M.* sp.1 is easily distinguishable from that species morphologically.

Although represented by a single sample, *M*. sp.2 clearly differed from other representatives of the *M. semirufus* complex based on both mitochondrial markers and our concatenated tree (Figures 1, 2, S1), although it shared a similar LwRh haplotype with most *M. semirufus* individuals (Figure S2). Interestingly, this sample was found almost at the southern border of *M. semirufus*’ distribution (Figure 3b). Although morphological characters suggest a potential differentiation between *M.* sp.2 and the rest of the *M. semirufus* complex, more samples from different locations are required in order to elucidate the status of this genetic lineage. We did collect a second colony in the same location as *M.* sp.2 (sample number 107 in Volcany Research Institute, see Table 1), but unfortunately, we could not amplify its DNA with any marker. Moreover, this sample was identified morphologically as *M. ebeninus*.

Another genetic lineage in Israel is represented by *M.* sp.3, which is clustered with samples of *M. minor* from Europe (Figure S3). Accordingly, these two entities keyed to the *M. minor* species complex based on morphological characters and formed a genetically distinct lineage in all our trees (Figures 1, 2, S1, S2, S3). We were able to collect only two individuals of this entity. In Vonshak & Ionescu-Hirsch (2009) *M. minor* is mentioned from the Jordan Valley and the Dead Sea area. It remains to be investigated whether it is rare in Israel. Clearly, more sampling is needed to confidently infer the two odd geographic sampling of *M*. sp.3 in Israel (Figure 3f) and determine how different it is from *M. minor*.

Lastly, *M.* sp.4 is another genetic lineage that we have recovered. *M.* sp.4 was retrieved as a distinct species in both the mitochondrial markers (Figures 1, S1) and our concatenated tree (Figure 2) but shared haplotypes with *M. ebeninus* and *“M. grandinidus”* on the nuclear marker (Figure S2). Its distribution appears mostly along the Central Jordan Valley (Figure 3d).

### Species Distribution Models (SDM)

The species distribution models performed well for all five examined lineages (Table S5). Variable contributions differed slightly among the five models, but soil type was the variable that contributed the most to explaining the model in four of the five lineages (Table 2). Other important contributing variables were annual temperature range (48.6% in *M.* sp.1, 20.8% in *M. ebeninus*, 11.9% in *M. semirufus*, all negatively correlated), annual precipitation amount (9.1% in *M. semirufus*, positively correlated), coarse fragments (9.1% in *M. ebeninus*, negatively correlated), and mixed/other trees (9.7% in *M. semirufus*, 8.3% in *M. ebeninus*, negatively correlated).

**Table 2:**
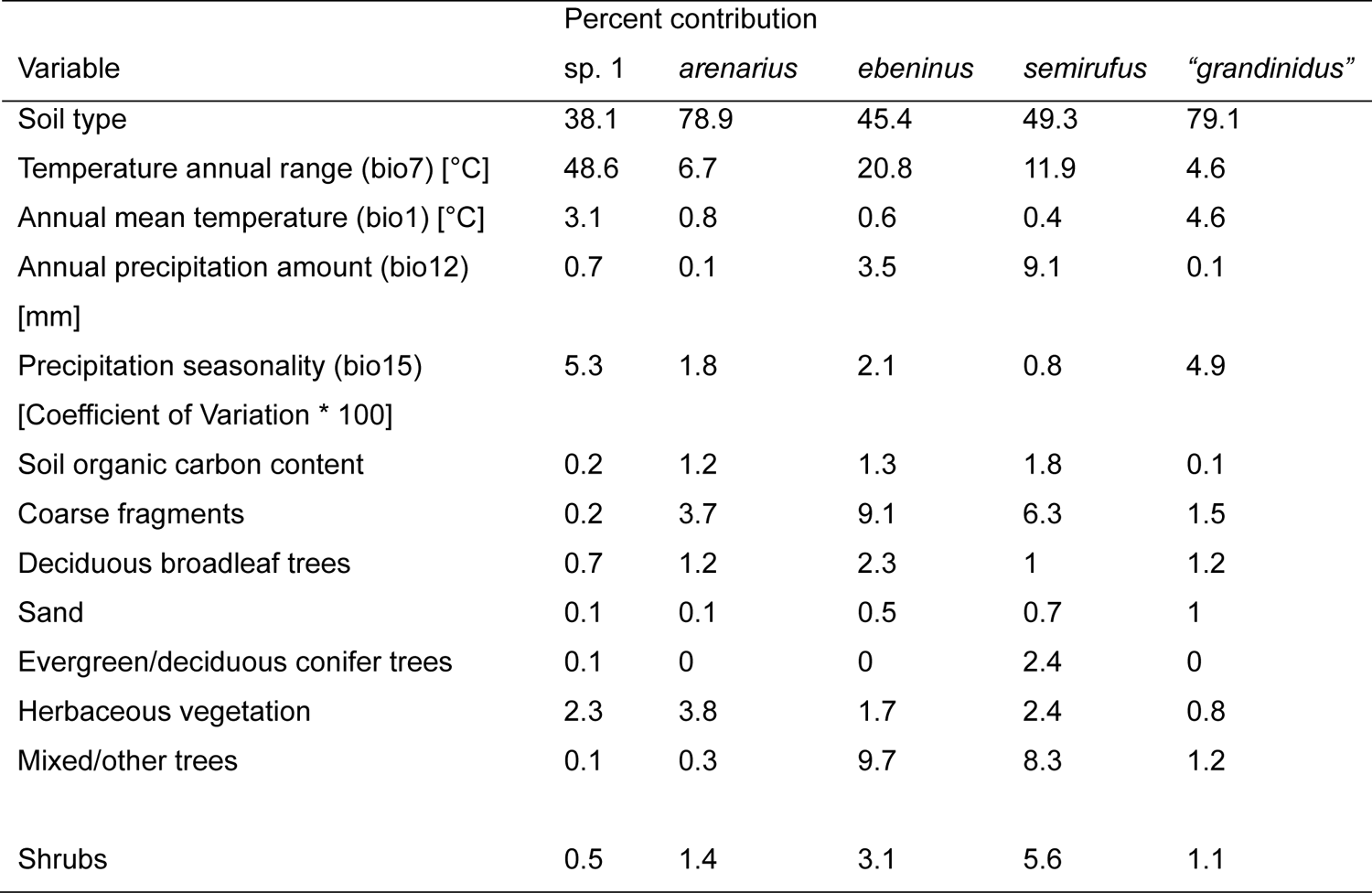
Variable contribution (%) to the model for five *Messor* species in Israel.

Soil type, which appeared to explain the probability of occurrences in most of the examined lineages, consisted of 23 distinct types (further subdivided into smaller categories; Itkin et al. 2018). Different soil types explained the probability of occurrence in the five lineages, with some overlap (Table 3). Additionally, prediction maps that were generated to identify areas with suitable habitats differed between the lineages (Figure S4). Habitats predicted to be suitable for “*M. grandinidus”* included major parts of the Negev and the Arava valley in the south of Israel (Figure S4c), whereas habitats predicted to be suitable for *M.* sp.1 were limited to the Central Coastal Plain (Figure S4e). *M. ebeninus* and *M. semirufus* both had relatively wide habitat predictions (Figure S4b, d), although *M. semirufus* seemed to be more limited to the central and northern parts of Israel. This finding is supported by our sampling of these species throughout Israel (Figure 3c, e). *M. arenarius* had a disjunct habitat prediction, consisting of the Central Negev and the Coastal Plain (Figure S4a). All of these findings are consistent with our genetic analysis.

**Table 3:** Soil types correlated with increased probability of occurrence of five *Messor* species in Israel. Asterisks indicate unique soil types per lineage.

| Lineage | Soils with increased occurrences |
| --- | --- |
| *"M. grandinidus"* | Terra rossas, brown rendzinas and pale rendzinas, brown lithosols and *loessial arid brown soils, brown lithosols and loessial serozems, *bare rocks and desert lithosols, Reg soils and coarse desert alluvium, loessial serozems, Sandy regosols and arid brown soils. |
| *M. semirufus* | Terra rossas, brown rendzinas and pale rendzinas, Grumusols, brown rendzinas and pale rendzinas, *basaltic protogrumusols, *basaltic brown grumusols and pale rendzinas, Hamra soils. |
| *M. ebeninus* | Terra rossas, brown rendzinas and pale rendzinas, Grumusols, *basaltic brown mediterranean soils and basaltic lithsols, brown rendzinas and pale rendzinas, pale rendzinas, sand dunes, Hamra soils, Reg soils and coarse desert alluvium, loessial serozems, Sandy regosols and arid brown soils. |
| *M. arenarius* | Sand dunes, Hamra soils, brown rendzinas and pale rendzinas, brown lithosols and loessial serozems, loessial serozems. |
| *M. sp.1* | Sand dunes, Hamra soils. |

## DISCUSSION

This study provides insights into the diversity, systematics and ecological preferences of *Messor* harvester ants of Israel, revealing the presence of at least 13 well-defined lineages, and highlighting the diversity and high level of endemism of *Messor* species in this ecologically diverse region. Species distribution models (SDM) that generated five of the Israeli lineages based on our sampling data-identified soil type and average annual temperature as the dominant factors explaining distribution patterns of *Messor* species in the country. The congruence between the phylogenetic clustering based on mitochondrial and nuclear genes and the predictions of ecological niche modelling lends strong support to the delineation of lineages here and demonstrates that species distribution modelling can serve as a valuable component for species delimitation.

Not all lineages that were supported by the phylogenetic analysis of the mitochondrial markers as distinct, were also supported by the nuclear marker (i.e., *M. ebeninus, M. semirufus, “M. grandinidus”, M. dentatus, M.* sp.2, and *M.* sp.4), which may attest to ongoing gene flow between these lineages via hybridization events, (Sukumaran & Knowles 2017). However, the clustering of all these six species into well-defined lineages in both mitochondrial markers despite their wide, overlapping distribution speaks against this scenario. Most likely, these results indicate relatively recent speciation of the relevant lineages (Goropashnaya et al. 2004), so that the shared most common nuclear (LwRh) alleles may stem from incomplete lineage sorting in this specific nuclear gene (Sullivan et al. 2002; Carstens & Knowles 2007). Hybridization and incomplete lineage sorting produce similar genetic signatures, and both are expected to occur in young species with incomplete reproductive barriers (Holder et al. 2001; Holland et al. 2008; Joly et al. 2009). In this case, the inclusion of faster evolving microsatellite markers may help to infer the occurrence of ongoing gene flow between these mitochondrial lineages (Jowers et al. 2014; Eyer et al. 2017). Similarly, behavioral assays testing mating preference and/or mating success between individuals belonging to different putative species can provide insights into the extent of reproductive isolation (Leppänen et al. 2016; Eyer et al. 2022).

Our sampling included individuals that key morphologically to *Messor maculifrons* Santschi 1927, *Messor hebraeus* Santschi 1927, and *Messor meridionalis* Andre 1883 */ Messor wasmanni* Krausse 1910, which are among the eight color variants of *M. semirufus* in Israel that were elevated to species level by Tohmé & Tohmé (1981). Given that these individuals did not form specific sub-grouping within *M. semirufus*, we concur with Kugler (1988) that they represent merely color variants of *M. semirufus* rather than distinct species. In addition to *M. semirufus*, Kugler (1988) recognized only *M. ebeninus* and *M. dentatus* as valid species. In accordance with our molecular analysis, we concur with Kugler on this observation as well. Colonies data points in Israel and the SDM analysis suggest that *M. ebeninus* and *M. semirufus* have relatively wide habitat preferences, resulting in wide distribution throughout Israel, although *M. semirufus* is more restricted to the central and northern parts of the country. This observation agrees with previous sampling of these species, with *M. ebeninus* recorded throughout Israel and *M. semirufus* occurring mainly in North Israel, the Judean Hills and along the Coastal Plain (Vonshak & Ionescu-Hirsch 2009).

*Messor grandinidus* inhabits North Africa, from Morocco to Libya (Stitz 1917; Santschi 1927; Finzi 1940) but genetic information did not exist for this species even in more recent works, i.e., Cagniant & Espadaler (1998) and Barech et al. (2020). The individuals in the “*M. grandinidus”* group that were sampled in this study are considered to be closely related to *M. grandinidus*. However, ants that were similar morphologically to those from the Negev were named *M. intermedius* in Arnol’di (1977); and other ants similar to those from the Arava Valley were identified as *M. ebeninus* or *M. meridionalis* in Collingwood & Agosti (1996). As we consider molecular discrimination to be most reliable, we suggest that differentiating between the 3 clades within the “M. grandinidus” lineage-that were recovered in Israel may serve as an important revenue for a further study. Sampel number 50, despite being only one individual-seems to be distinct from the other 3 clades, as the two mitochondrial haplotype network analyses depict, which supports a fourth potential clade within this lineage (Figures 1, S1). The single individual belonging to this group that was recorded from the Coastal Plain (sample number 81_1; see Figure 3c for the distribution) was collected at a site where Mount Carmel reaches the Mediterranean Sea. This geographic region harbors different soils and is inhabited by different ant species than the rest of Israel’s Coastal Plain (Eyer et al. 2017; Ionescu-Hirsch & Eyer 2016), which may explain this odd sampling point.

*Messor* sp.1 clustered close to *M. aegyptiacus fellah* but differs from it morphologically, and our SDM analysis confirmed that the distribution of *M.* sp.1 is limited to the central Coastal Plain (Figures 3f, h, S4e), unlike *M. aegyptiacus fellah-* strongly suggesting that this is an undescribed species. More sampling of *M. aegyptiacus fellah* are needed to elucidate its relationship with *M.* sp.1, especially considering the fact that both species appear to be locally endemic (Vonshak & Ionescu-Hirsch 2009).

*M.* sp.3 clearly differentiated from other *Messor* lineages in all genetic markers, but in contrast to the morphologically unique *M.* sp.1, its morphological distinctiveness from *M. minor* remains unclear. Given the results of the molecular analysis, we re-examined the only two minute (small sized) specimens available for *M*. sp.3 and found morphological similarities to specimens from Israel and Saudi Arabia that have previously been identified as *M. minor*.

Other erroneous identifications revealed in this study include one sample (number 54) that was identified morphologically as “*M. grandinidus”* but was genetically assigned to *M. ebeninus,* and similarly-several individuals that were identified as *M. semirufus* were genetically assigned to *M. ebeninus* (samples number 42, 43, 52, and 53). However, these erroneous identifications were consistent through all of our molecular analysis. Moreover, morphologically identifying *Messor* species with the tool keys that we used currently-sometimes led to the identification of individuals from a single colony-as belonging to several different species, especially in the northern part of Israel. One concrete example is the case of colony number 81 from Ramat Ha’nadiv, where we morphologically identified two individuals - one belonging to *“M. grandinidus”* (81_1) and the other to *M. semirufus* (81_2). Surprisingly, in this case the identification was supported by our two mitochondrial and concatenated analyses (Figures 1, 2, S1). These erroneous or baffling identification efforts suggest that keys to the *Messor* species in this region should be revised. A similar situation was reported for the *Messor structor* complex in Europe, where the main diagnostic morphological characters (width of the metasternal process and length of the scape) varied among individuals within colonies (Schlick-Steiner et al. 2006). Consequently, a thorough molecular, morphological, and ecological analysis revealed that *M. structor* actually comprised five cryptic species (Steiner et al. 2018). Lastly, one sample in our study (number 10) could not be identified morphologically, and thus remained *M.* sp. (without a number). Nonetheless, the sample was assigned genetically to the “*M. grandinidus”* lineage.

Predicted distributions produced by the species distribution models were largely congruent with the observed ranges of the genetically inferred lineages in Israel. For most lineages, soil type and average annual temperature were the variables explaining most of the variance. Preferences for certain soil types were shared by all lineages, but some types were unique to three of the five lineages (asterisks in Table 3) Soil types used by a single lineage may represent environments with low competition from other *Messor* species (Solida et al. 2010, 2011a, b; Steinberger et al. 1992), although they may still be inhabited by other ant genera. Interestingly, *M. arenarius* and *M. sp.1* were not associated with unique soil types but were found to inhabit fewer soil types in specific habitats, and preferentially nest in sand dunes along the coast or in sandy areas of the Northern Negev, suggesting a rather specialist nature.

The influence of abiotic factors such as soil properties and topography on distribution, survival, and establishment has been reported in congeneric *Messor barbarus* Linnaeus 1767 colonies (Baraibar et al. 2011; Enzmann & Nonacs 2010; Johnson 1998; Wiernasz & Cole 1995, Crist & Wiens 1996). Toughness, texture, and moisture of the topsoil affected the ants’ ability to excavate and build nest chambers (Boulton et al. 2005; Enzmann & Nonacs 2010; Johnson 1998; Wiernasz & Cole 1995). In *M. barbarus*, only one percent of founding colonies succeeded in establishing (Gordon & Kulig 1996) and the survival of these colonies remained low between 2-5 years thereafter (Gordon 1995; Johnson 2001), emphasizing the importance of establishing colonies in a suitable soil type. Similarly, in three species of *Pogonomyrmex* harvester ants, nest digging rate and nest depth were positively associated with the degree of claustrality during colony foundation, namely, whether founding queens rely on their internal reserves (claustral) or on foraging (not claustral) to raise their first brood of workers (Enzmann & Nonacs 2010). Overall, these observations highlight that variation among harvester ant species in their ability to excavate soil may promote certain founding strategies or excavation behaviors depending on the soil type (Johnson 1992).

In addition to influencing colony foundation and nest construction, different soil types may also be associated with certain flora communities, which produce a variety of seed types and sizes. In turn, this may influence species diversity of harvester ants as many are known to preferentially forage on specific seed types (Johnson 2000; Belchior et al. 2012; Kugler & Hincapié 1983; Kugler 1984; Pirk et al. 2009; Pol et al. 2011; Pol & Lopez-de-Casenave 2004; MacMahon et al. 2000; Solida et al. 2010, 2011a; Rissing 1986, 1988; Wilby & Shachak 2000). Moreover, in *Messor*, mean worker size is often associated with the size of harvested seeds (and other plant material) (Rissing 1981; Waser 1998; Heredia & Detrain 2005; Azcárate et al. 2005; Willott et al. 2000). *Messor* species with different mean worker sizes are expected to forage on seeds of different sizes, and therefore may experience lower interspecific competition (Rissing & Pollock 1984; Heredia & Detrain 2005; Plowes et al. 2012; Vorster et al. 1991). The high variability in foraging strategies exhibited by different *Messor* species can also reduce the ecological competition for a resource, allowing for the co-occurrence of several species at a given locality (Plowes et al. 2012; Saar et al. 2018; Pol et al. 2022). This means that soil type affects not only colony foundation success, excavation behavior, and nest structure of *Messor* species, but may shape their distribution patterns also by influencing the environment and resources available to them. The use of soil type as an explanatory variable is therefore particularly relevant when modelling the distribution of species in this ant group. As soil properties also influence the survival and distribution of many other ant species (e.g., Boulton et al. 2005; Kaspari & Weiser 2007; Ríos-Casanova et al. 2006; Schoereder & DaSilva 2008; Costa-Milanez et al. 2017), using soil type variables is likely to offer meaningful support when modelling the distribution of other ant species, as has been demonstrated for *Cataglyphis* spp. Foerster 1850 in Israel (Eyer et al. 2017, 2018).

In summary, we report here of the presence of newly discovered species and species complexes in the *Messor* harvester ants in Israel, some of which are possibly rare and endemic. We also raise *M. bodenheimeri* and *M. ratus* from subspecies to valid species. Although we were unable to sample all the *Messor* taxa reported from Israel by Vonshak & Ionescu-Hirsch (2009), our morphological, genetic, and ecological analyses point to necessary development of accurate keys for local *Messor* species and call for further investigation of these taxa in this ecologically diverse region.

## ACKNOWLEDGMENTS

This study was supported by a VATAT (The Planning and Budgeting Committee, the Israeli Council for Higher Education) post-doctoral fellowship (for MS) for research at the Steinhardt Museum of Natural History, Tel-Aviv University. MS was also supported by a Vaadia-BARD Postdoctoral Fellowship, Award No. FI-595-19 from the United States-Israel Binational Agricultural Research and Development Fund. We are grateful to A. Blumenfeld for his comments on the manuscript.

## AUTHORS CONTRIBUTIONS

MS conceived and planned the research, managed collaboration between the coauthors, collected data, performed the molecular work and wrote the manuscript. PAE collected data, performed the molecular analyses and wrote the manuscript. TMC performed the niche modelling analyses and wrote on the manuscript. AIH provided taxonomic input and wrote the manuscript. RD and ND funded the research, provided conceptual advice and wrote the manuscript.

## SUPPORTING INFORMATION

**Table S1:** Genes and primers used in this study.

**Table S2:** PCR amplification conditions for markers used in this study.

**Table S3:** Environmental variables used in the niche modelling analyses.

**Table S4:** Pearson’s r correlations between niche environmental variables in the modelling analyses.

**Figure S1:** (*a*) Network of mitochondrial CytB haplotypes of *Messor* harvester ants in Israel. Colors correspond to lineages. Branch lengths indicate the number of mutations between haplotypes; circle sizes are proportional to the number of haplotypes observed in the dataset; (*b*) Bayesian tree of mitochondrial CytB haplotypes *Messor* in Israel based on the mitochondrial CytB marker. Branch support values are Bayesian posterior probabilities. Numbers 1-3 in squares represent postulated clades within the *M. semirufus* and “*M. grandinidus*” lineages.

**Figure S2:** Network of nuclear LwRh haplotypes of *Messor* harvester ants in Israel. Branch lengths indicate the number of mutations between haplotypes; circle sizes are proportional to the number of haplotypes observed in the dataset. Dashed lines represent heterozygous haplotypes co-occurring in the same individual, their weight proportional to the number of heterozygous individuals. Colors correspond to mitochondrial lineages of origin.

**Figure S3:** Bayesian tree of *Messor* harvester ants in Israel (colored in correspondence with lineages in Figures 1, 2, 3, S1, S2) and reference species from GenBank (colored in gray), based on the mitochondrial COI marker. Branch support values are Bayesian posterior probabilities.

**Figure S4:** Maps of predicted suitable habitats for five *Messor* lineages in Israel, according to the niche modelling analysis. Colors represent probabilities, ranging from low (blue) to high (red).

