## Supplementary material for "Systematics of harvester ants (*Messor*) in Israel based on integrated morphological, genetic, and ecological data": Table S1, Table S2, Table S3, Table S4, Table S5, Figure S1, Figure S2, Figure S3, Figure S4

**No. of Tables & Figures: 5 & 4**

**Tables**

**Table S1:** Genes and markers details

| **mtDNA/nDNA** | **Gene** | **Marker** | **Sequences (5' to 3')** | **References** |
| --- | --- | --- | --- | --- |
| mtDNA | COI | Messorfor2 PatMessor | F: CGGCTCTCTAGGAATAATTTATGC  R: AATCTATTGCACTTTTCTGCCA | Steiner et al. 2011 |
| mtDNA | CytB | CB1  CB2 | F: TATGTACTACCATGAGGACAAATATC  R: ATTACACCTCCTAATTTATTAGGAAT | Chiotis et al. 2000; Moreau 2008 |
| nDNA | LwRh | LR143F | F: GACAAAGTKCCACCRGARATGCT | Ward & Downie 2005; Brady et al. 2006 |

**Table S2:** PCR amplification conditions per gene

| **Gene** | **Initialization** | | **35-40 thermal cycles**  **Denaturation Annealing Elongation** | | | | | | **F. Elongation** | |
| --- | --- | --- | --- | --- | --- | --- | --- | --- | --- | --- |
|  | Min | Temp | Min | Temp | Min | Temp | Min | Temp | Min | Temp |
| *COI* | 3 | 94 | 0.5 | 94 | 0.5 | 48 | 1 | 72 | 10 | 72 |
| *CytB* | 2 | 94 | 1 | 94 | 1 | 48 | 1 | 72 | 10 | 72 |
| *LwRh* | 1 | 95 | 0.5 | 95 | 0.5 | 58 | 1.5 | 72 | 3 | 72 |

Min = Minutes, Temp = Temperature at ° C, F. Elongation = Final elongation.

**Table S3:** Details of environmental variables resources used in this study

| **Variable type** | **Variable** | **Definition [unit]** | **Original resolution [cell size]** | **Reference** | **Used in the final model** |
| --- | --- | --- | --- | --- | --- |
| Soil | Carbon | Soil organic carbon content (stock) [‰ (g kg−1)] | 0.002 | Hengl et al., 2014, 2017 | Yes |
|  | Coarse fragments | Coarse fragments [volumetric %] |  |  | Yes |
|  | Sand | Sand [weight %] |  |  | Yes |
|  | Clay | Clay [weight %] |  |  |  |
|  | Bulk density | Bulk density [kg m^-3^] of the fine earth fraction (< 2 mm) |  |  |  |
|  | Silt | Silt [weight %] |  |  |  |
|  | Cation | Cation-exchange capacity [cmol + /kg] of the fine earth fraction |  |  |  |
|  | pH | Soil pH in H2O and KCl solution |  |  |  |
|  | Soils | Soil types | 646.208 | Itkin et al., 2018 | Yes |
| Land cover | Deciduous | Deciduous broadleaf trees [consensus prevalence in percentage] | 0.008 | Tuanmu and Jetz., 2014 | Yes |
|  | Evergreen | Evergreen/deciduous needleleaf trees [consensus prevalence in percentage] |  |  | Yes |
|  | Herbaceous | Herbaceous vegetation [consensus prevalence in percentage] |  |  | Yes |
|  | Mixed | Mixed/other trees [consensus prevalence in percentage] |  |  | Yes |
|  | Shrubs | Shrubs [consensus prevalence in percentage] |  |  | Yes |
| Climate | Bio1 | Annual mean temperature [°C] | 0.008 | Karger et al., 2017 | Yes |
|  | Bio2 | Mean diurnal range [°C] |  |  |  |
|  | Bio3 | Isothermality [Bio2/Bio7 * 100] |  |  |  |
|  | Bio4 | Temperature seasonality [SD * 100] |  |  |  |
|  | Bio5 | Maximum temperature of warmest month [°C] |  |  |  |
|  | Bio6 | Minimum temperature of coldest month [°C] |  |  |  |
|  | Bio7 | Temperature annual range [°C] |  |  | Yes |
|  | Bio8 | Mean temperature of wettest quarter [°C] |  |  |  |
|  | Bio9 | Mean temperature of driest quarter [°C] |  |  |  |
|  | Bio10 | Mean temperature of warmest quarter [°C] |  |  |  |
|  | Bio11 | Mean temperature of coldest quarter [°C] |  |  |  |
|  | Bio12 | Annual precipitation amount [mm] |  |  | Yes |
|  | Bio13 | Precipitation of wettest month [mm] |  |  |  |
|  | Bio14 | Precipitation of driest month [mm] |  |  |  |
|  | Bio15 | Precipitation seasonality [Coefficient of Variation * 100] |  |  | Yes |
|  | Bio16 | Precipitation of wettest quarter [mm] |  |  |  |
|  | Bio17 | Precipitation of driest quarter [mm] |  |  |  |
|  | Bio18 | Precipitation of warmest quarter [mm] |  |  |  |
|  | Bio19 | Precipitation of coldest quarter [mm] |  |  |  |

**Table S4:** Pearson’s r correlations between environmental variables used in this study. *P*-values are indicated above the diagonal, and Pearson’s r values are indicated below the diagonal.

|  | **Carbon** | **Sand** | **Coarse fragments** | **Shrubs** | **Herbaceous** | **Mixed** | **Evergreen** | **Deciduous** | **Bio1** | **Bio7** | **Bio12** | **Bio15** | **Soils** |
| --- | --- | --- | --- | --- | --- | --- | --- | --- | --- | --- | --- | --- | --- |
| Carbon |  | <0.001 | <0.001 | <0.001 | <0.001 | <0.001 | <0.001 | <0.001 | <0.001 | <0.001 | <0.001 | <0.001 | <0.001 |
| Sand | 0.52 |  | <0.001 | <0.001 | <0.001 | <0.001 | 0.459 | 0.006 | <0.001 | <0.001 | <0.001 | <0.001 | <0.001 |
| Coarse fragments | 0.43 | 0.80 |  | <0.001 | <0.001 | <0.001 | 0.319 | 0.013 | <0.001 | <0.001 | <0.001 | <0.001 | <0.001 |
| Shrubs | 0.40 | 0.04 | 0.07 |  | <0.001 | <0.001 | <0.001 | <0.001 | <0.001 | <0.001 | <0.001 | 0.001 | <0.001 |
| Herbaceous | 0.26 | 0.01 | -0.01 | 0.16 |  | <0.001 | 0.385 | <0.001 | <0.001 | <0.001 | <0.001 | <0.001 | <0.001 |
| Mixed | 0.21 | -0.01 | -0.03 | 0.12 | 0.03 |  | <0.001 | <0.001 | <0.001 | <0.001 | <0.001 | <0.001 | <0.001 |
| Evergreen | 0.06 | 0.00 | 0.00 | 0.02 | 0.00 | 0.23 |  | <0.001 | 0.001 | <0.001 | <0.001 | <0.001 | <0.001 |
| Deciduous | 0.16 | -0.01 | -0.01 | 0.16 | 0.08 | 0.22 | 0.16 |  | <0.001 | <0.001 | <0.001 | <0.001 | <0.001 |
| Bio1 | -0.29 | 0.15 | -0.11 | -0.26 | -0.09 | -0.07 | -0.01 | -0.06 |  | <0.001 | <0.001 | <0.001 | <0.001 |
| Bio7 | -0.53 | 0.23 | 0.55 | -0.24 | -0.31 | -0.21 | -0.06 | -0.20 | -0.06 |  | <0.001 | <0.001 | <0.001 |
| Bio12 | 0.73 | -0.47 | -0.51 | 0.47 | 0.33 | 0.24 | 0.06 | 0.21 | -0.20 | -0.74 |  | <0.001 | <0.001 |
| Bio15 | 0.03 | 0.08 | 0.05 | 0.01 | -0.15 | -0.06 | -0.02 | -0.09 | 0.09 | 0.13 | 0.06 |  | <0.001 |
| Soils | -0.58 | 0.65 | 0.46 | -0.44 | -0.17 | -0.21 | -0.05 | -0.18 | 0.33 | 0.62 | -0.80 | 0.24 |  |

**Table S5:** Average Area Under the Curve (i.e., AUC) values for each of the five examined clades.

| **Lineage** | **Average AUC** | **Standard deviation** |
| --- | --- | --- |
| *“M. grandinidus”* | 0.953 | 0.019 |
| *M. semirufus* | 0.986 | 0.004 |
| *M. ebeninus* | 0.988 | 0.005 |
| *M. arenarius, M. ratus* | 0.989 | 0.005 |
| *M.* sp.1 | 0.998 | 0.002 |

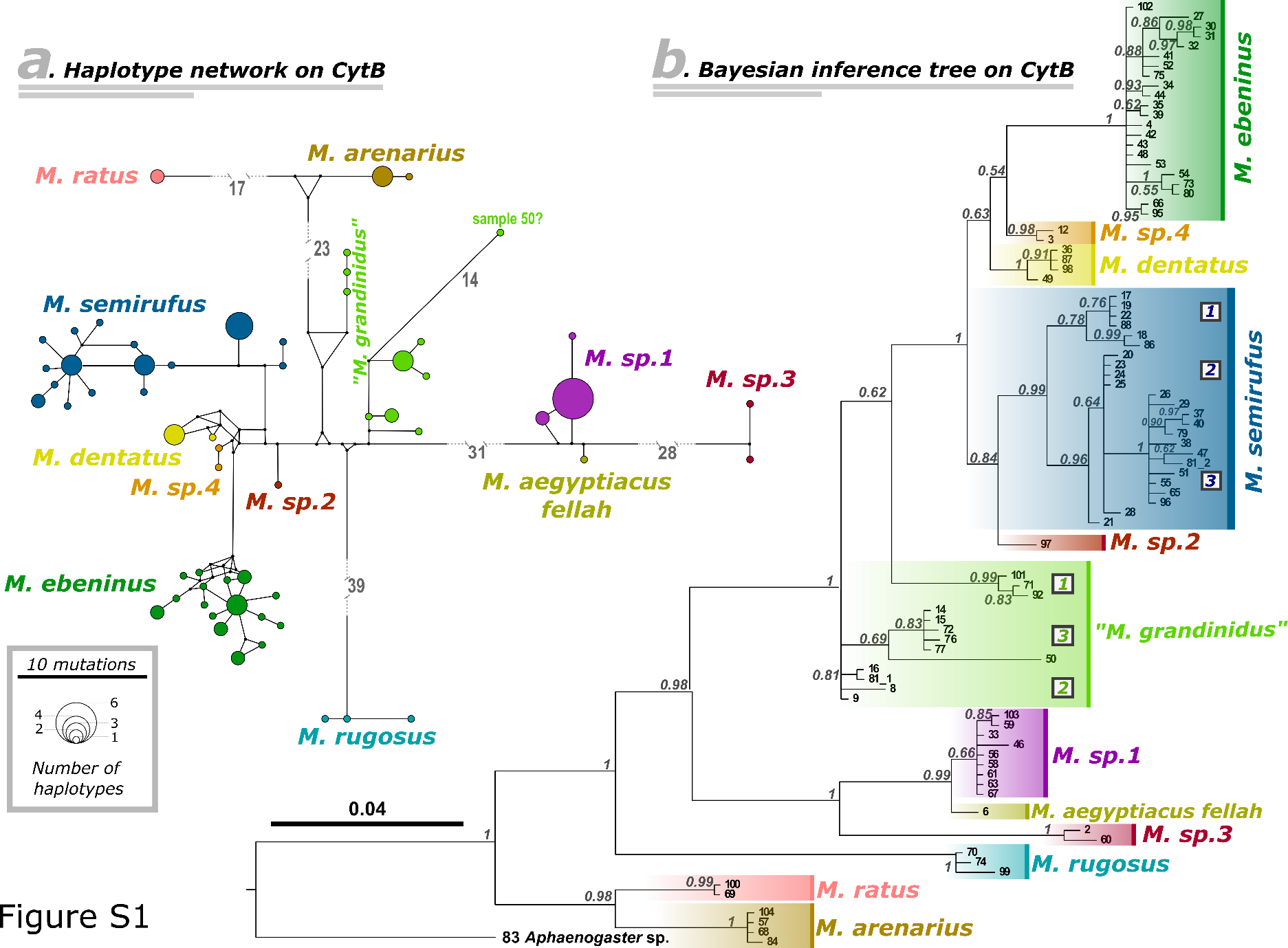

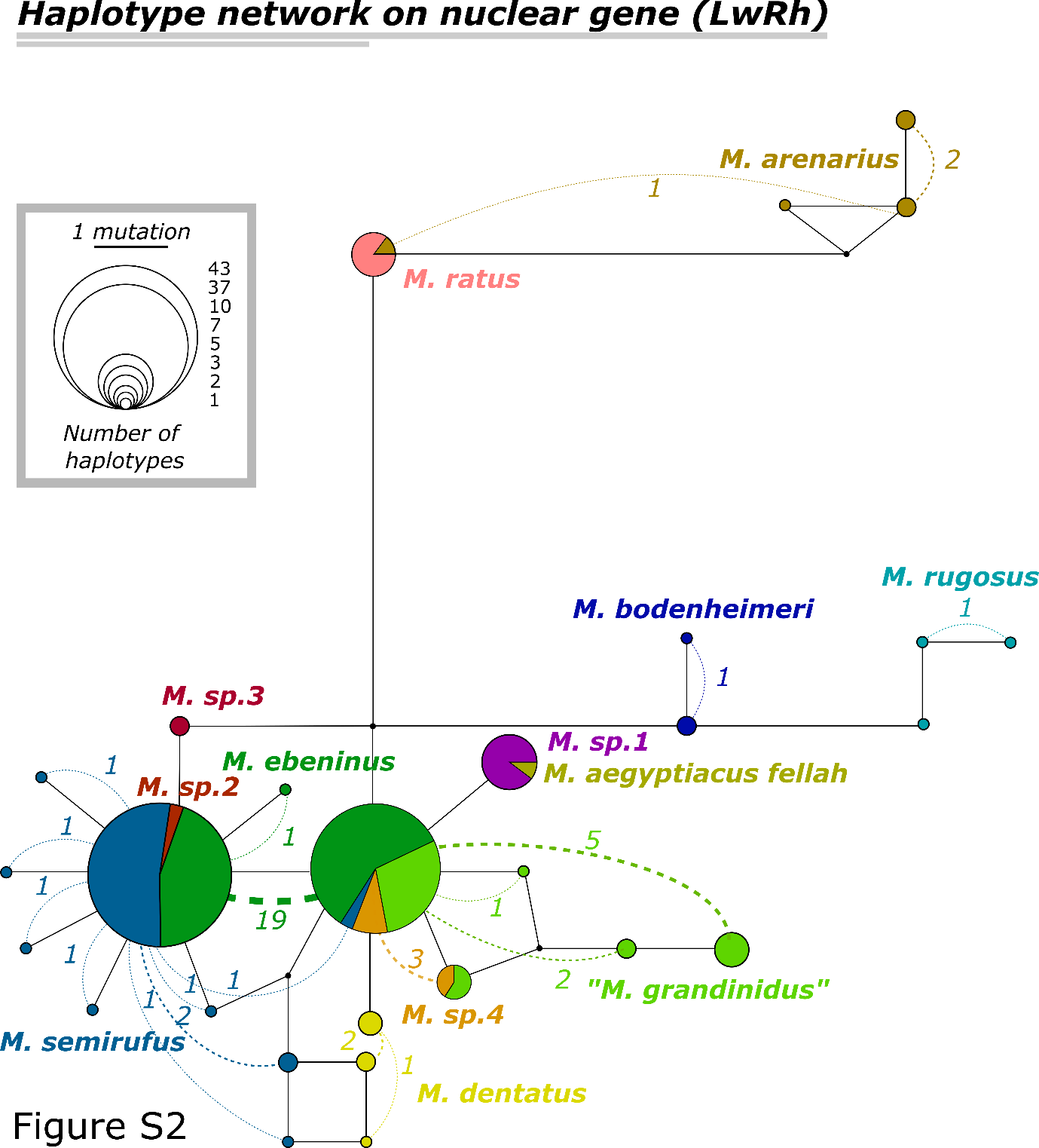

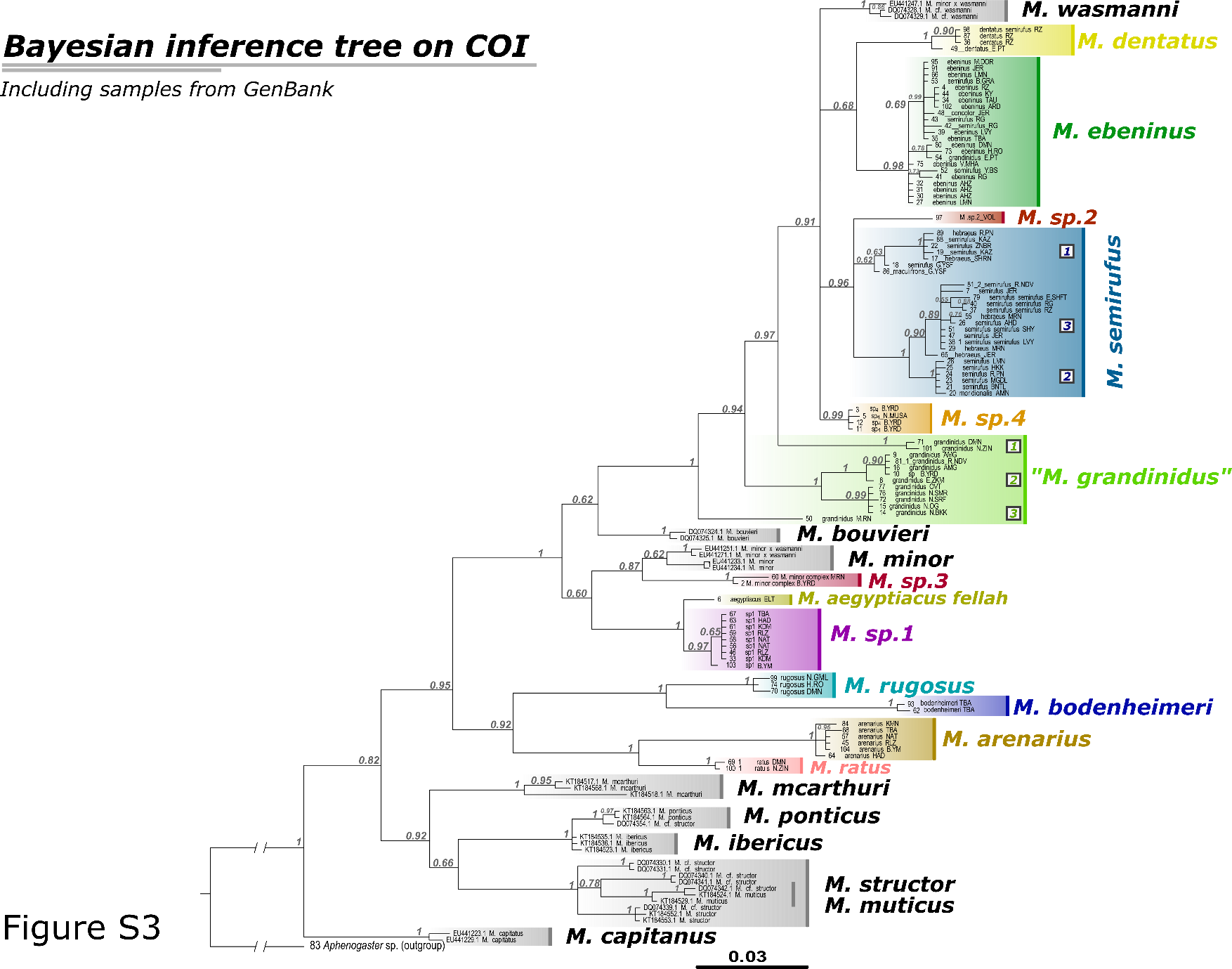

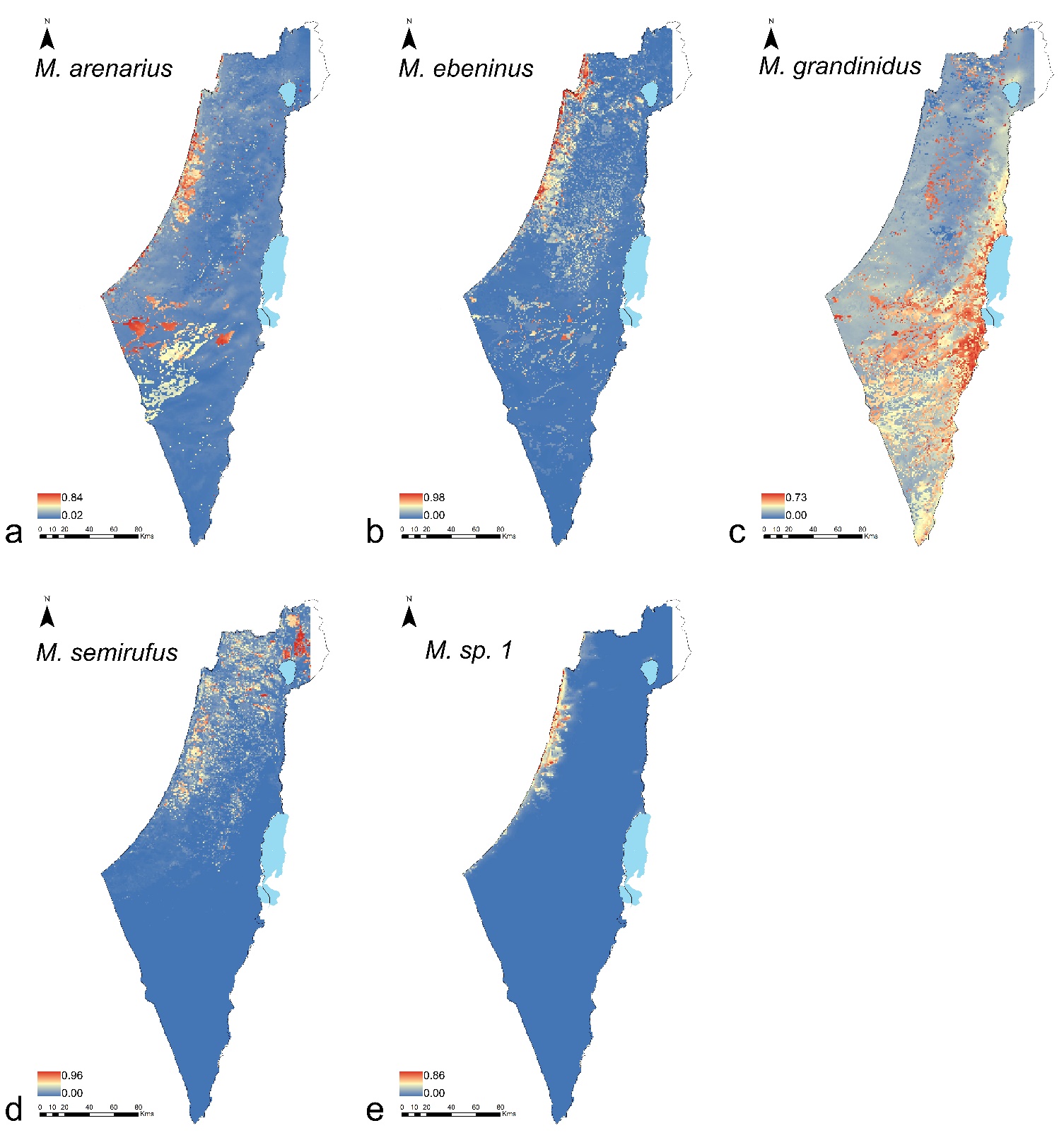

**Figure S4**
